# Genomic signatures of long-term selection in the Langhill dairy herd: the power of a five-decade experiment for sustainable dairy cattle breeding

**DOI:** 10.64898/2026.11.06.764806

**Authors:** Rajiv Sharma, Mike Coffey, Georgios Banos, Raphael Mrode, Bingjie Li, Ian Archibald, Eileen Wall, Wayne Powell

## Abstract

**Background:** Selective breeding has transformed dairy cattle productivity, but the genomic consequences of long-term directional selection — what it costs in diversity and where in the genome it acts — are hard to quantify because contemporaneous control populations are rare. The Langhill experiment, established in 1973, is one of the world’s longest-running dairy cattle selection trials with a designed control line kept under common management. We used this resource population to ask what five decades of bull selection have cost in genetic diversity, where in the genome selection has acted, and whether complementary scans converge on the loci under selection.

**Results:** We genotyped archived blood samples from 1,126 cows of two genetic lines: a Select line sired by bulls in the top decile of UK genetic merit for milk fat-plus-protein yield, and a Control line sired by bulls of average merit. The Select line showed the expected gains in every yield trait. Inbreeding accumulated at the same annual rate in both lines, and effective population size, estimated from that rate, was approximately 33. Linkage disequilibrium was higher in the Select line at every marker distance from 2 to 300 kb. Autozygosity was 24.7% higher in the Select line than in the Control line (0.078 versus 0.063), a difference robust across four calling parameter sets and two independent callers. The excess was confined to runs of homozygosity longer than 2 Mb; shorter runs did not differ between lines, confirming a shared founder base and placing all divergence after the lines were split. It did not arise from a numerically narrower sire pool: the Select line used more sires (79 versus 72), each with fewer daughters, yet its cows were three times more related to one another. Twenty-eight regions across thirteen chromosomes showed strong differentiation; intersecting five scans identified a four-method signal on chromosome 21 spanning the SERPINA gene cluster and sixteen three-method regions, including chromosome 3, with haplotype and diversity statistics agreeing on the line carrying the signal in fourteen of seventeen. Autozygosity itself did not localise to any region.

**Conclusions:** Selection left two genomic footprints: diffuse, recent inbreeding accumulated through the narrow ancestry of the elite sire pool, and localised differentiation at specific loci. Runs of homozygosity index only the first process; the second is captured by frequency- and haplotype-based differentiation scans. Because the cost of inbreeding is incurred through ancestry rather than the number of bulls used, mating schemes that constrain the rate of co-ancestry would be preferable to simply managing the number of sires selected and used.

## Introduction

Livestock agriculture supplies billions of people with food and underpins rural livelihoods worldwide^1,2^, yet it faces mounting pressure to reduce its environmental footprint whilst continuing to raise efficiency^3,4^. Conscious selection and breeding have been central tools for improving productivity, and breeding will be equally central to lowering emissions intensity and supporting adaptation to a changing climate^5,6^. Understanding how directional selection reshapes the genome over time is therefore of direct practical relevance to the design of future breeding programmes^7,8^. Yet the genomic cost of this success — how much standing variation directional selection consumes, and where in the genome it is spent — has remained difficult to quantify, because in commercial populations selection is inextricably confounded with drift, sire migration and a continually shifting management environment.

Long-term selection experiments offer a uniquely controlled window onto this process^9,10^. Here we exploit the Langhill dairy herd experiment, established in Scotland in 1973 and one of the world’s longest-running dairy cattle selection trials^11^. From a single Holstein-Friesian base population, two lines have been maintained: a Control line, repeatedly sired by bulls of UK-average genetic merit, and a Select line, sired by bulls in the top decile of UK merit for fat-plus-protein yield^12,13^. From 1988 a high-versus low-concentrate feeding contrast was nested within each line^13^. Because the Select line tracks the genetic trajectory of the elite national herd while the Control line approximates the contemporaneous average UK population, the experiment provides an unusually direct estimate of the cumulative genomic impact of sire selection. This is what distinguishes Langhill from comparisons of separate breeds, or of a single unselected line against contemporary commercial cows: both lines descend from one base population and have been bred in parallel under identical husbandry for five decades, with feeding system the only deliberately varied environmental axis. Breed background, era and environment are thereby largely controlled, so that genomic divergence between the lines can be attributed to sire choice with a confidence rarely attainable outside an experimental cross.

Genotyping of archived blood samples allows this trajectory to be read retrospectively. Although selection signatures have been mapped in several Holstein populations, these studies have largely contrasted breeds or set one unselected line against contemporary cows^14–16^, leaving the genome-wide consequences of a controlled, like-for-like selection contrast unresolved. Reading the Langhill lines genomically therefore allows changes in diversity, homozygosity and locus-specific differentiation to be attributed directly to selection, and tests whether complementary statistics converge on the same regions. We ask how sustained selection has altered phenotypic means, population structure, linkage disequilibrium, nucleotide diversity and homozygosity, and we identify genomic regions bearing signatures of divergent selection. We interpret the findings against comparable selection-signature studies in Holstein and other dairy populations^14,17–19^, notably a U.S. Holstein line unselected since 1964 in which selection for milk production left footprints overlapping fertility and immunity genes^14,15^, and discuss their implications for maintaining diversity in climate-resilient dairy breeding. We test whether five decades of directional selection have measurably eroded genome-wide diversity, elevated homozygosity and produced convergent signatures at specific loci, and we read these against the fertility and health responses documented in the same animals. The result is a controlled accounting of what genetic gain has cost in genomic terms — a template for weighing productivity against the erosion of the genomic diversity on which future, climate-resilient breeding depends.

The present study addresses two primary objectives regarding the genomic consequences of long-term directional sire selection. First, we quantify the impact on genome-wide genetic diversity—measured through nucleotide diversity, linkage disequilibrium (LD), and autozygosity—and resolve these patterns across a 50-year birth-cohort trajectory. Second, we use high-density genomic data to link the line contrast to genomic regions associated with the traits under selection, testing whether complementary selection scans converge on consistent genomic loci. As context for both, we first confirm the phenotypic response to selection in the genotyped cows. To achieve these objectives, we integrate genome-wide diversity statistics and length-resolved decomposition of runs of homozygosity (ROH) with five complementary scans for selection signatures.

## Results

### Phenotypic impact of selection

Across the recorded period, the Select line significantly exceeded the Control line for the selection-objective traits — total daily milk yield, and daily protein and fat yield — and for the morning, midday and evening milking components (Figure 1A–F; all P < 0.001). Select cows were also heavier (612.9 vs 590.7 kg; P = 1.1 × 10⁻⁶) and were recorded at a slightly later stage of lactation (week 25.5 vs 23.3; P = 0.039). Consistent with the established correlated response in this herd, fertility (calving interval) and body condition score declined under selection until the introduction of the UK fertility index and have since partially recovered. Direct selection in the Select line was applied on milk fat-plus-protein yield alone; the recovery in fertility is therefore most plausibly an indirect consequence of the bull pool from which Select sires were drawn having been pre-selected by breeding companies on fertility once the UK fertility index became available^20^.

**Figure 1.**
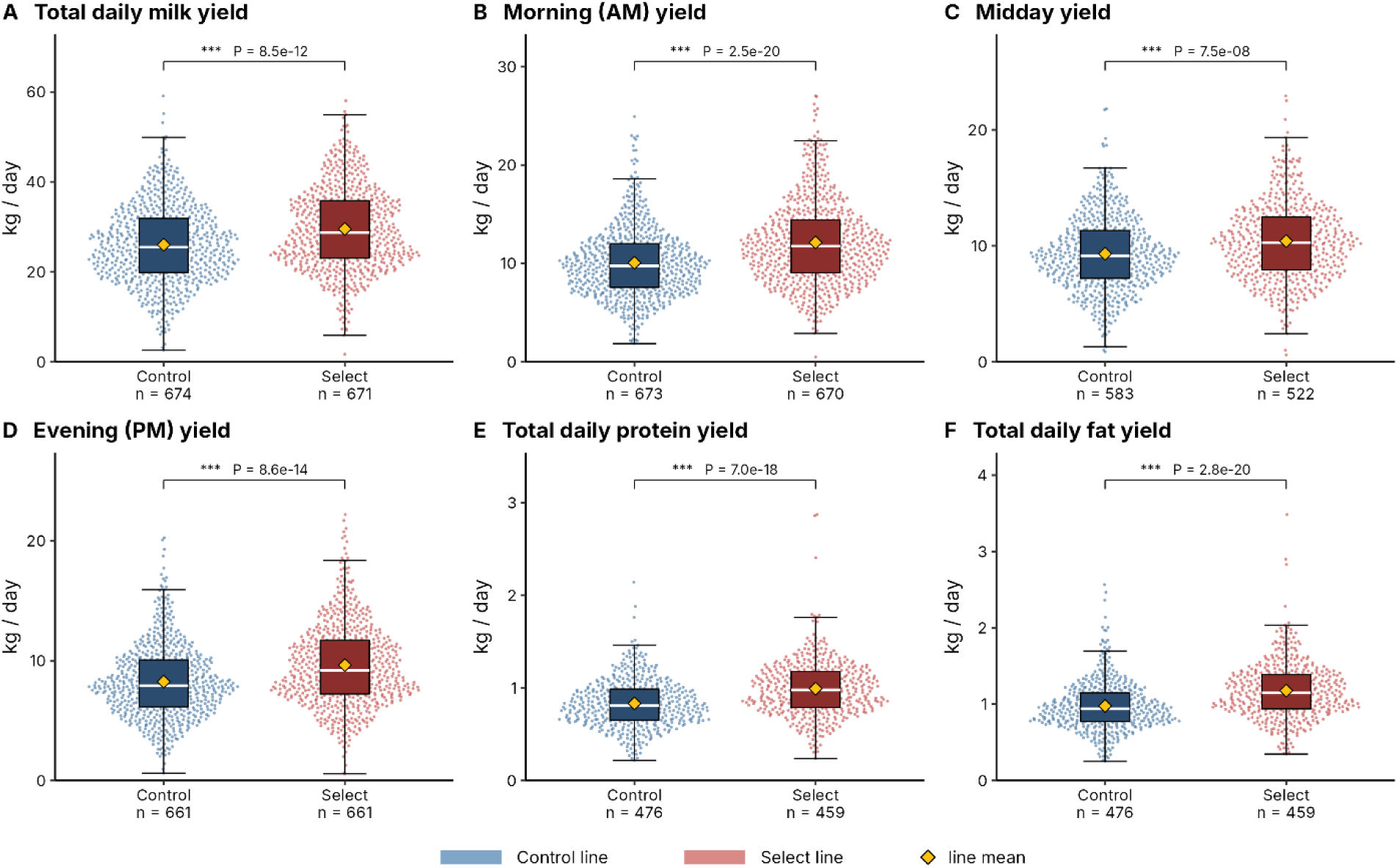
Phenotypic divergence between Langhill selection lines. Beeswarm plots with embedded box plots for (A) total daily milk yield, (B) morning (AM) yield, (C) midday yield, (D) evening (PM) yield, (E) total daily protein yield and (F) total daily fat yield. Control blue, Select red; yellow diamonds are line means, white lines medians; points are jittered in proportion to local density. Zero values were treated as missing and excluded, so n differs by trait (shown per panel). P values are two-sided Mann–Whitney tests; *** P < 0.001.

### Population structure and linkage disequilibrium

Principal component (PC) analysis separated the two lines along PC1 and PC2 (2.50% and 1.29% of variance explained, respectively) (Figure 2A), computed from 17,852 LD-pruned autosomal single nucleotide polymorphism (SNP) markers. The 53 cows that clustered with the opposite line (Figure 2A,B) were concentrated in the earliest cohorts: 41 of the 47 Select cows assigned to the Control-like cluster were born before 2000, when the lines were least differentiated, and the remainder are consistent with the occasional parentage errors documented in the herd records. The modest variance on the leading axes is expected within a single breed split into two recently diverged lines, where genome-wide differentiation is low and distributed across many axes. Line membership was taken from the herd breeding records throughout; model-based clustering was used to validate that assignment and, for the differentiation and haplotype scans only, to exclude the 53 discordant cows (see Methods). At an ancestry-coefficient threshold of Q > 0.6, fastSTRUCTURE assigned every genotyped cow to a cluster and recovered the recorded line in 1,073 of 1,126 cases (95.3%; Figure 2B).

**Figure 2.**
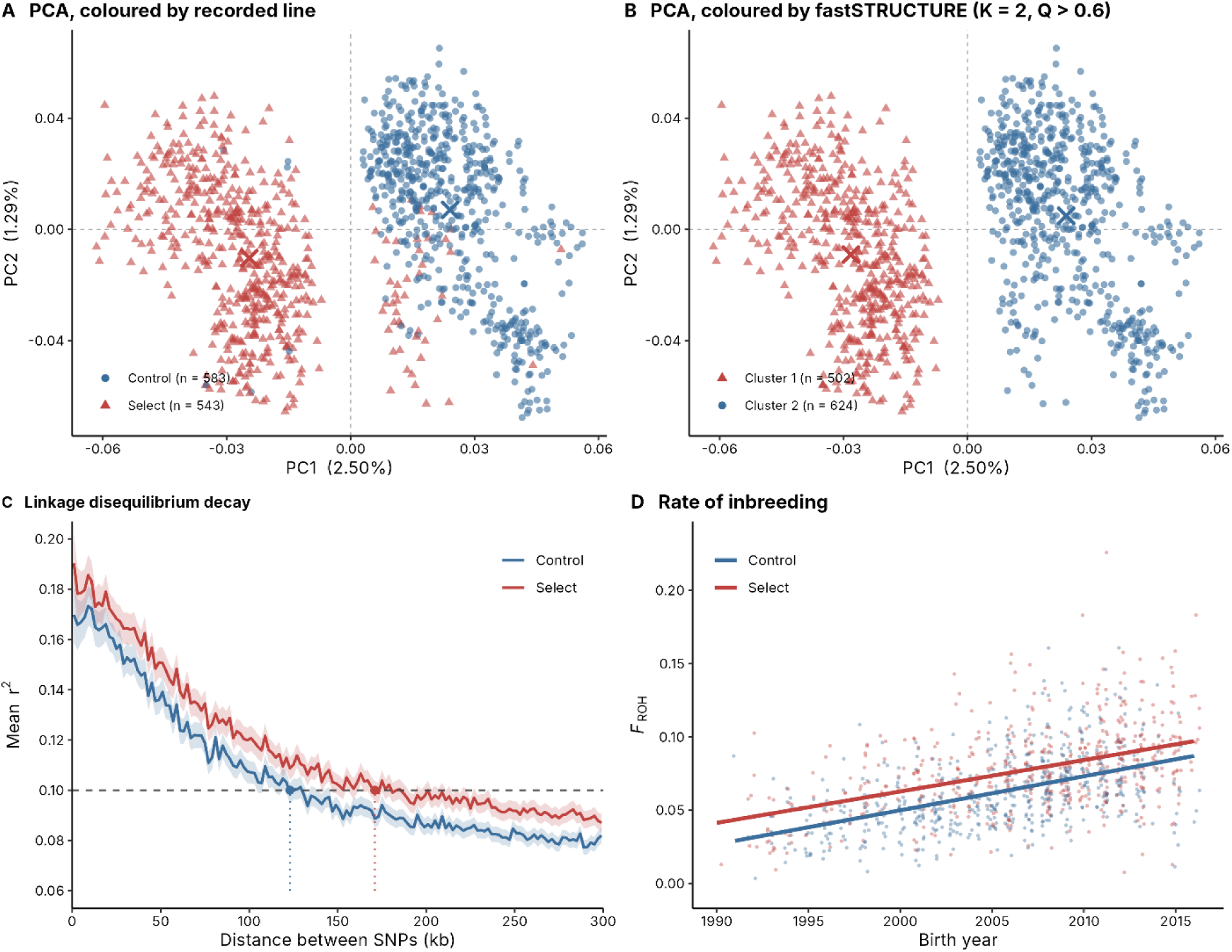
Population structure, linkage disequilibrium and effective population size. (A) Principal components 1 and 2 from 17,852 LD-pruned autosomal markers in 1,126 cows, coloured by line as recorded in the herd breeding records; PC1 explains 2.50% and PC2 1.29% of variance. Crosses mark line centroids. (B) The same principal components, coloured instead by fastSTRUCTURE assignment at K = 2 using an ancestry-coefficient threshold of Q > 0.6. Every genotyped cow is assigned at this threshold, and the assignment agrees with the breeding records for 1,073 of 1,126 cows (95.3%); of the 53 discordant cows, 47 are recorded as Select but cluster with Control (41 of them born before 2000) and 6 the reverse. Panels A and B show the same points under two independent labellings: model-based clustering recovers the recorded design without using it, which is why line membership was taken from the records throughout. (C) Linkage disequilibrium decay: mean r² between marker pairs against physical distance, in 2-kb bins weighted by pair count and computed separately per line; shading is the 95% confidence interval. The horizontal dashed line marks r² = 0.10, which the Control line first reaches at 123 kb and the Select line at 171 kb — 48 kb further; dotted droplines and points mark those crossings. Linkage disequilibrium is higher in the Select line at every distance from 2 to 300 kb, by more than twice the width of the confidence intervals throughout. (D) Individual F_ROH against birth year (points jittered horizontally for legibility) with least-squares fits per line. The rate of inbreeding does not differ between lines (year × line interaction P = 0.49); both share a slope of +0.0022 F_ROH per year, while the Select line sits +0.012 higher throughout (P = 1.6 × 10⁻¹⁵), an offset already present in cows born in 1990. Using the generation intervals measured from the pedigree (Control 8.2, Select 5.4, pooled 6.8 years), ΔF per generation is 0.0189 (Control) and 0.0117 (Select), giving effective population sizes of 26 and 43 respectively, and 33 from the pooled common slope, which is the estimate reported throughout. Control blue circles, Select red triangles throughout.

Linkage disequilibrium was higher in the Select line than in the Control line at every marker distance from 2 to 300 kb, and by more than twice the width of the 95% confidence intervals throughout (Figure 2C; Table S3). Mean r² first fell to 0.10 at 171 kb in the Select line but at 123 kb in the Control line, and at 100 kb was 0.120 versus 0.107. Slower decay indicates longer-range haplotype correlation — longer shared blocks — in the Select line, the expected consequence of stronger directional selection and associated drift/hitchhiking rather than a difference in recombination rate. Consistent with this, genomic relatedness among cows was substantially higher within the Select line (mean KING kinship 0.0040 vs 0.0013; *P* = 2.3 × 10⁻¹⁷⁶), despite the Select line having close pairs no more often (1.46% vs 1.61% of pairs at kinship > 0.125): the Select line is more uniformly related through distant shared ancestry rather than through recent close matings.

### Genome-wide nucleotide diversity

Nucleotide diversity (π), representing the average number of pairwise nucleotide differences per site, was modestly lower in the Select line (median 4.415 vs 4.420 and mean 4.234 vs 4.265 × 10⁻⁵ per bp; lower on 24 of 29 autosomes; Figure 3A; per-chromosome profiles in Figure S1). Because 93% of 10-kb windows on this array contain a single marker, adjacent windows are not independent and window-level significance tests are not reported; the difference is small relative to that background. Temporal change in diversity is instead captured by autozygosity, which is measured per animal and requires no window-based approach (Figure 2D). At the level of birth cohorts, which avoids window pseudoreplication, mean π declined across cohorts in the Select line (−0.017 × 10⁻⁵ per cohort step; 95% CI −0.026 to −0.008, chromosome bootstrap) but not in the Control line (−0.004; −0.012 to 0.004; Table S1).

**Figure 3.**
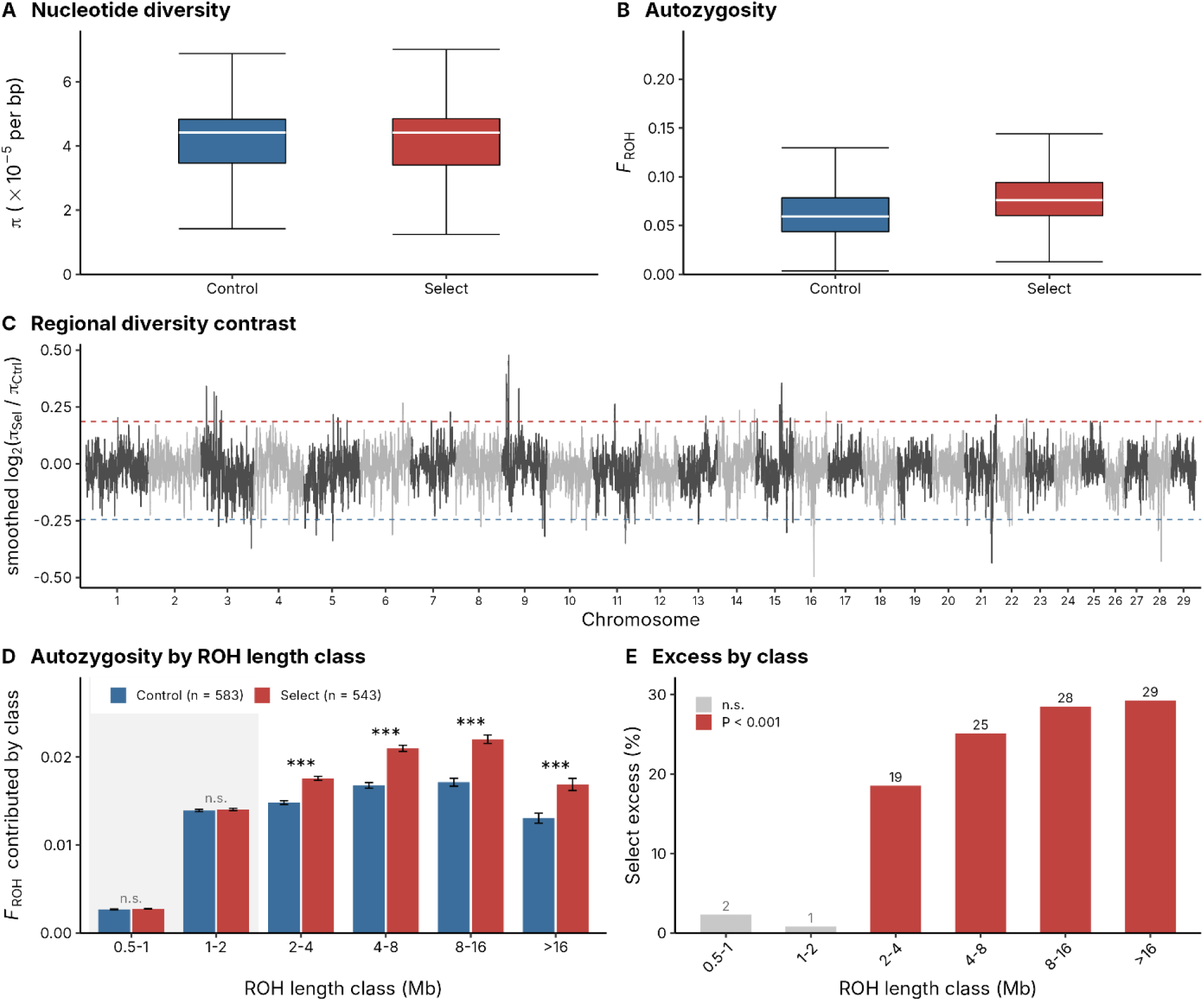
Genome-wide diversity and autozygosity. (A) Nucleotide diversity (π) per 10-kb window by line; the view is clipped at the upper whisker because π has a long right tail (maximum 44 ×10⁻⁵ against an interquartile range of 3.4–4.8). (B) F_ROH by line: mean 0.0779 in the Select line versus 0.0625 in the Control line, a 24.7% excess (Mann–Whitney P = 9.4 × 10⁻²²). (C) Smoothed log₂(π_Select/π_Control) across autosomes 1–29 (25-window rolling mean); dashed lines mark the 0.5% tails (−0.244 and +0.186), and windows beyond them are carried forward to the cross-method intersection. (D) F_ROH partitioned by ROH length class (ROH called at 30 SNPs / 500 kb); bars are means, error bars standard errors; the shaded panel marks the short classes. Classes below 2 Mb do not differ between lines (0.5–1 Mb P = 0.29; 1–2 Mb P = 0.45, both n.s.) whereas every class above 2 Mb does (all P ≤ 2.1 × 10⁻⁵), locating the Select line’s excess autozygosity in recent rather than founder ancestry. (E) Select-line excess per class: +2.3%, +0.8%, +18.5%, +25.1%, +28.5% and +29.3% respectively, grey where not significant. Control blue, Select red throughout; n = 583 Control and 543 Select cows.

Regional contrasts of the smoothed π ratio identify discrete windows in the 0.5% tails (Figure 3C), which are carried forward to the cross-method intersection. Because per-window π ratios are dominated by sampling noise on this marker density, they are used only in combination with the other scans, never alone.

### Autozygosity

Autozygosity was markedly higher in the Select line: mean F_ROH 0.0779 versus 0.0625, a 24.7% excess (Mann–Whitney P = 9.4 × 10⁻²²; Figure 3B). The difference was insensitive to ROH-calling parameters, the absolute excess varying only between +0.0147 and +0.0158 across four parameter sets spanning an eight-fold range of minimum length and two independent callers (Table 3).

Decomposing F_ROH by segment length separates ancient from recent autozygosity (ROH called at 30 SNPs / 500 kb so that 0.5–1 Mb segments are resolved; Figure 3D,E; Table 5). ROH shorter than 2 Mb, which reflect founder haplotypes shared across the base population, did not differ between lines (0.5–1 Mb: +2.3%, P = 0.29; 1–2 Mb: +0.8%, *P* = 0.45). Every class above 2 Mb differed, and the excess grew with length (2–4 Mb: +18.5%; 4–8 Mb: +25.1%; 8–16 Mb: +28.5%; >16 Mb: +29.3%; all P ≤ 2.1 × 10⁻⁵), such that 99% of the excess lay in ROH exceeding 2 Mb. Equal short-ROH burdens confirm that both lines retain the same founder autozygosity and that all divergence has accumulated since the lines were split. The excess arose principally through more numerous segments (37.0 vs 31.1 per cow, +19%, *P* = 8.3 × 10⁻²³) rather than longer ones (5,139 vs 4,887 kb, +5%, *P* = 1.9 × 10⁻⁶). Per-cow segment counts and mean segment lengths are given in Table S2.

This additional autozygosity was not concentrated at specific loci. Because genome-wide F_ROH is itself higher in the Select line, the expected odds of any locus falling within a long ROH is already elevated in that line irrespective of any locus-specific process (genome-wide median odds ratio 1.25). Testing every autosomal marker against a null that preserves each cow’s genome-wide ROH burden and permutes line within strata of that burden, the most differentiated locus in the genome (BTA5 ∼8.4 Mb; odds ratio 6.46) was no more extreme than permutation routinely produces (*P* = 0.74). The excess autozygosity of the Select line is therefore genome-wide rather than localised.

Autozygosity accumulated at the same rate in both lines. Regressing individual F_ROH on birth year, the year × line interaction was not significant (*P* = 0.49): the two lines share a slope of +0.0022 F_ROH per year while the Select line sits +0.012 higher throughout (P = 1.6 × 10⁻¹⁵), an offset already present in cows born in 1990. Because Select cows have a later mean birth year, this year-adjusted offset (+0.012, equivalent to a 19% excess at the Control mean) is smaller than the unadjusted difference of +0.0154 (24.7%). The divergence in autozygosity was therefore established early and has been maintained rather than enlarged. Based on the shared rate and using the generation interval measured from the pedigree (6.8 years; Methods), the rate of inbreeding per generation (Δ*F*) is 0.0151. This corresponds to an effective population size (*N_e_*) of approximately 33 (33–38 depending on how the generation interval is summarised, and 42 under the 5.4-year interval typical of dairy populations; Figure 2D), a value consistent with the restricted census size of this closed experimental herd. Because the Control line was sired by older, progeny-proven bulls, its generation interval is longer (8.2 versus 5.4 years), so a similar annual rate translates into more inbreeding per generation in the Control line (ΔF = 0.0189 versus 0.0117, each from the line’s own slope and generation interval; Ne ≈ 26 versus 43). The higher per-generation estimate for the Select line should therefore not be read as greater diversity: per year, autozygosity accumulates at the same rate in both lines, and the lines differ in its level, which was established early and has been maintained. Linkage disequilibrium (LD) fails to provide an independent estimate of *N_e_* in this context. Because the generations sampled by an LD-based estimator scale as *t* = 1/(2*c*), the 300-kb window analyzed only reaches approximately 167 generations into the past—a temporal depth that far exceeds the 50-year span of this experiment.

### Genetic differentiation

Genome-wide differentiation was measured with FST (between-line share of total genetic variance) and ZFST (Z-standardised 10-SNP rolling-mean form, used for the extreme tail). Mean per-SNP FST was low at 0.023, as expected within a pure breed^15,17^. The ZFST scan identified twenty-eight strongly differentiated regions (ZFST > 5, merged within 1 Mb) across thirteen chromosomes (Figure 4; Table 1). Chromosome 3 carried the most regions (n = 7), followed by chromosome 9 (n = 6); chromosomes 8 and 9 carried the highest peaks (ZFST = 7.61 at chr9 32.7 Mb; 7.71 at chr8 41.2 Mb). A further notable region was chromosome 9 at ∼97.7 Mb (peak rolling FST = 0.125), which coincides with the only chromosome-9 π reduction in the Select line. In total 96 markers exceeded ZFST > 5, delimiting the 28 regions reported here. The ZFST, XP-EHH and Rsb scans used the 1,073 cows whose recorded line agreed with genomic clustering (see Methods). Repeating the ZFST scan on all 1,126 cows lowered mean FST to 0.020 but left the regions essentially unchanged: 26 of the 28 again exceeded ZFST > 5, the remaining two reached 4.97 and 4.99, and the leading peaks were identical.

**Figure 4.**
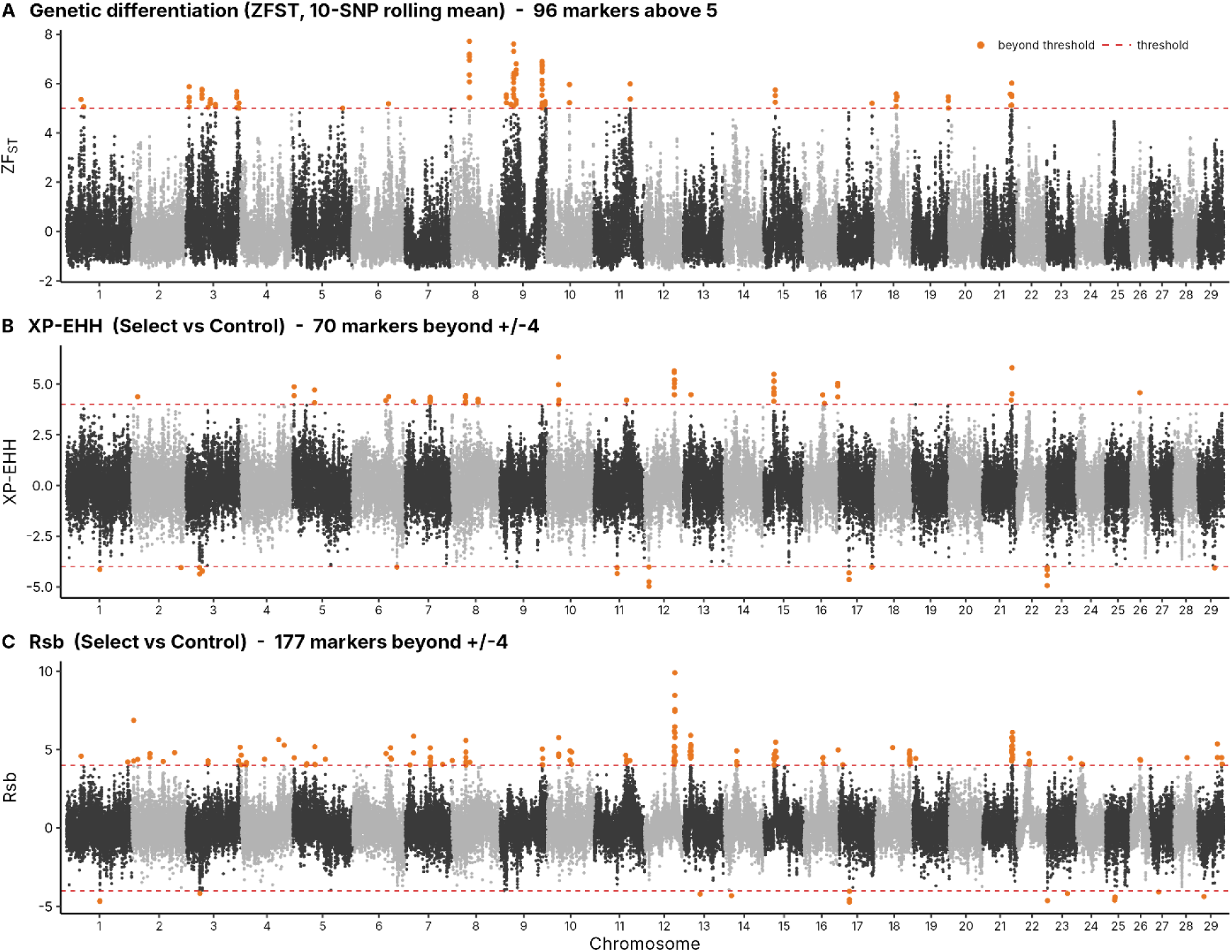
Genetic differentiation and haplotype-based selection signals. Genome-wide scans across autosomes 1–29; alternating shading distinguishes chromosomes and orange marks the markers beyond the threshold (dashed). (A) ZFST, Z-standardised 10-marker rolling mean of per-marker Weir–Cockerham FST; 96 markers exceed 5, delimiting 28 regions across thirteen chromosomes; peaks BTA8 41.2 Mb (Z = 7.71) and BTA9 32.7 Mb (Z = 7.61). (B) XP-EHH (Select vs Control); 70 markers exceed |4|, delimiting 32 regions (21 with positive scores). (C) Rsb (Select vs Control); 177 markers exceed |4|, delimiting 73 regions (63 with positive scores); strongest signal BTA12 ∼71.6 Mb (Rsb = 9.90). Positive values indicate extended haplotype homozygosity in the Select line.

**Table 1.**
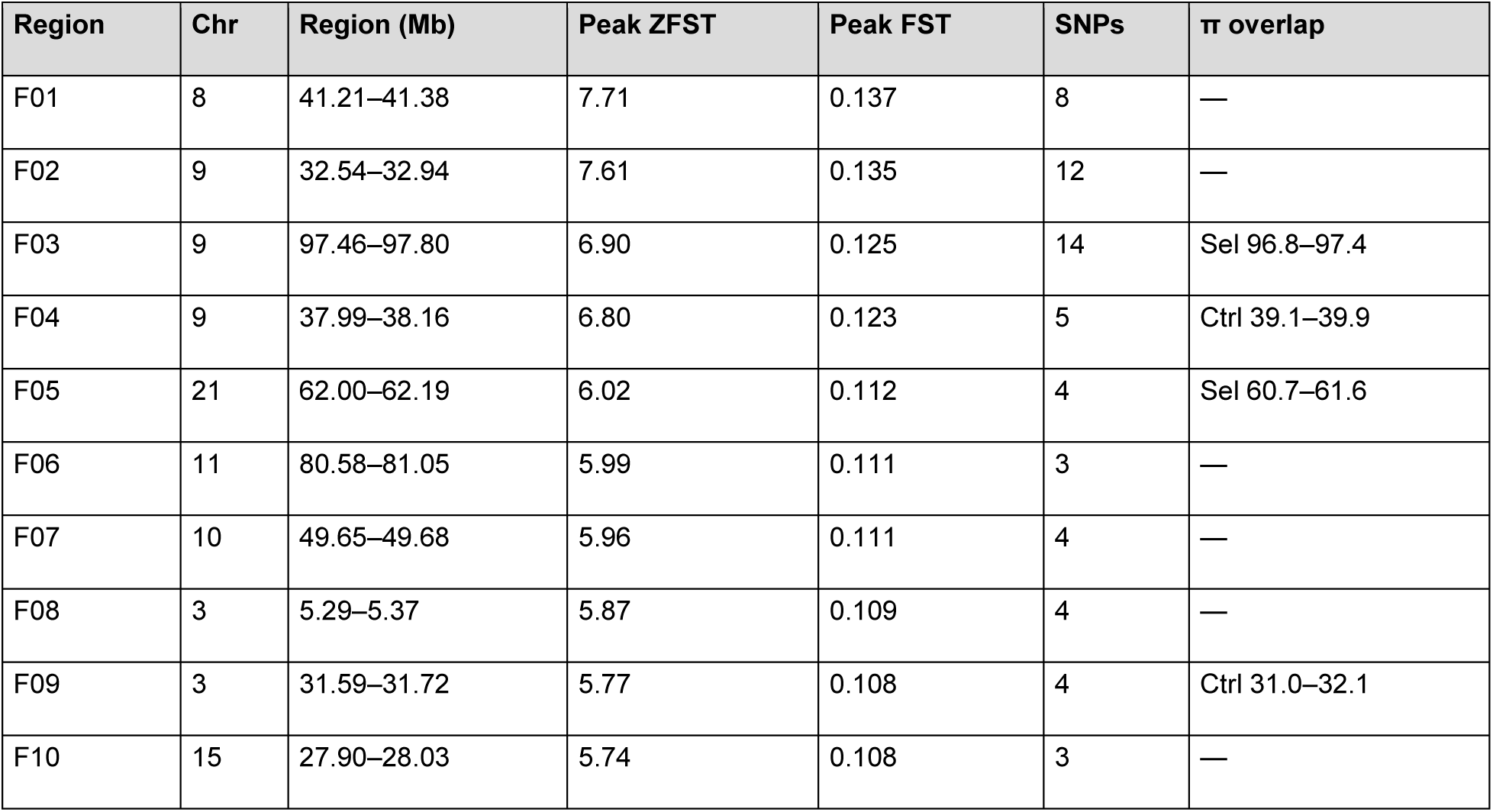
Top 10 of 28 differentiated regions (ZFST > 5), ranked by peak ZFST; full list in Table S4. Coordinates ARS- UCD1.2. Genome-wide mean per-SNP FST = 0.023.

### Haplotype-based signals and cross-method convergence

Cross-population extended haplotype homozygosity (XP-EHH) and the Rsb statistic^21^ — sensitive to near-fixed sweeps — identified 32 and 73 regions, respectively (70 and 177 markers with |score| > 4, merged within 1 Mb; 21 and 63 of them with extended haplotype homozygosity in the Select line; Figure 4B,C; Table S5). Merging regions from all five scans within 1 Mb gave 132 candidate regions, of which 45 were detected by ≥2 methods and 17 by ≥3 (Table 2; Table S6; Figure 5). One region, chromosome 21 (59.2–62.2 Mb), was recovered by all four statistics (ZFST, XP-EHH, Rsb and the diversity contrast) and is the strongest convergent selection candidate in the experiment; chromosome 3 (31.4–32.2 Mb) was recovered by three (ZFST, XP-EHH and the diversity contrast). Haplotype and diversity statistics, computed separately, agreed on the line carrying the signal in 14 of the 17 regions: ten, including chromosome 21, showed extended haplotype homozygosity and reduced π in the Select line, and four, including chromosome 3 at 31.4 Mb, showed the reverse, consistent with drift or selection within the Control line (Table 2). Separately, the chr9 ∼97.7 Mb differentiated region (Table 1, F03) co-locates with a Select-line π reduction, and the chr16 47.8–48.8 Mb region is shared by XP-EHH, Rsb and π. As expected, the two highest FST peaks (chr8 41.2 Mb; chr9 32.7 Mb) are haplotype-frequency signals not mirrored in the sparse-array π. The strongest Rsb signal, and the largest single value in any scan, was on BTA12 at ∼71.6 Mb (Rsb = 9.90), where the three highest markers cluster. The second-strongest XP-EHH peak (BTA21, 62.0 Mb; XP-EHH = 5.80) falls inside the BTA21 four-method convergent interval, providing haplotype-based support for that region in addition to its ZFST and π signals. The single highest XP-EHH value (BTA10, 22.3 Mb) was less stable in resampling checks than the BTA21 and BTA3 signals and is not interpreted further (Supplementary Note S1).

**Figure 5.**
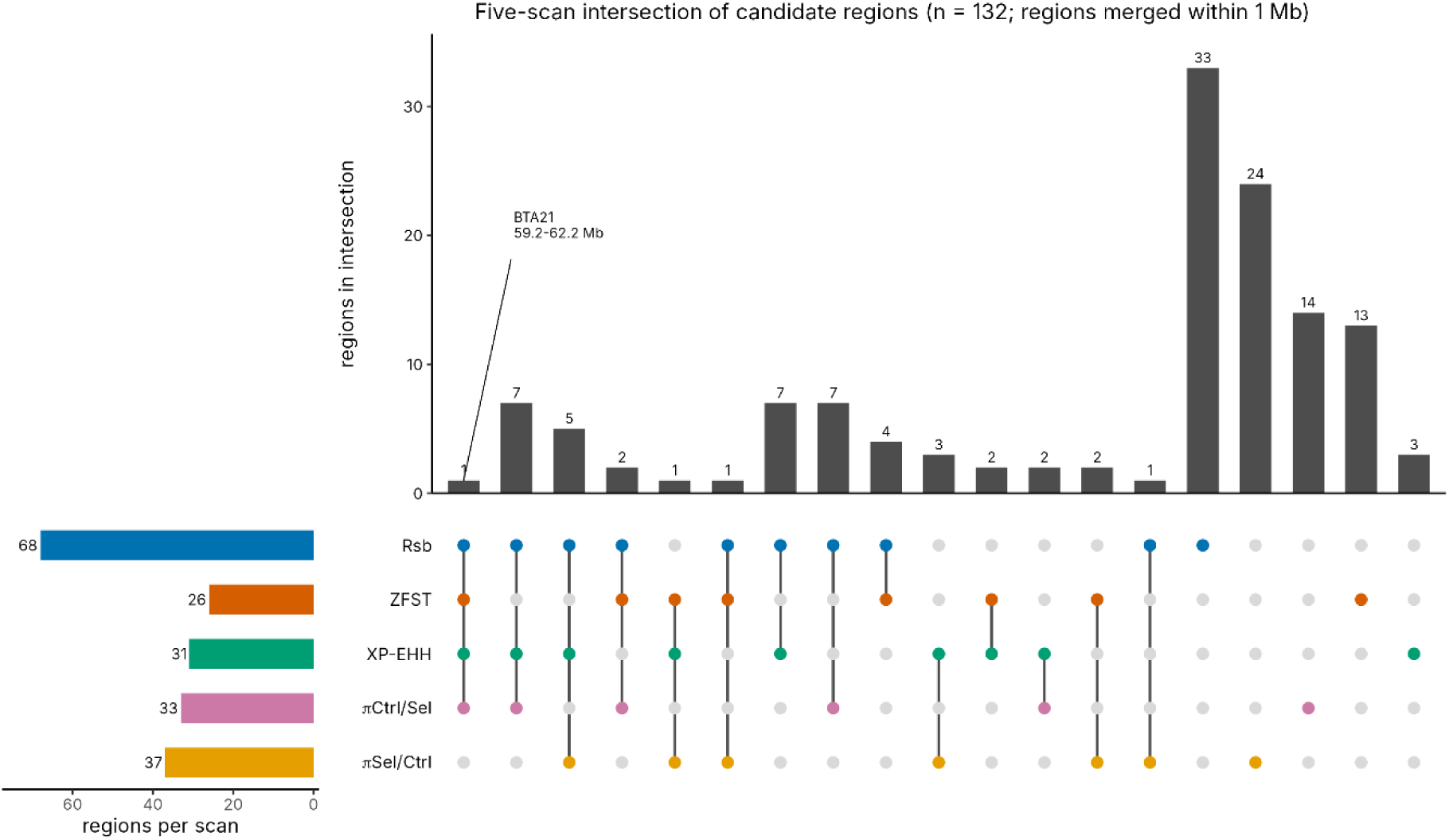
Cross-method convergence. UpSet plot of the 132 candidate regions obtained by merging regions from the five scans within 1 Mb; one region (BTA21, 59.2–62.2 Mb) is supported by four scans and 16 by three (Table 2; Table S6). The bars on the left give the number of candidate regions containing a region from each scan (ZFST 26, XP-EHH 31, Rsb 68, π Control > Select 33, π Select > Control 37); these are lower than the per-scan region counts in the text (28, 32 and 73 for ZFST, XP-EHH and Rsb) because regions from different scans within 1 Mb of one another are merged into a single candidate region.

**Table 2.** The 17 candidate regions detected by ≥3 of the five scans (one by four), regions merged within 1 Mb; ±250 kb flanks used for annotation. Line: the line showing extended haplotype homozygosity (sign of XP- EHH/Rsb) and, after the semicolon, reduced π. Genes: Ensembl release 104 protein-coding genes within ±250 kb (up to six named genes shown; full lists in Table S8). QTL: number of Animal QTLdb entries within ±250 kb and the three leading QTL classes (Table S9). Positions are ARS-UCD1.2.

| Region | Chr | Region (Mb) | ±250 kb | Methods | Line; genes; QTL<sup>22,23</sup> |
|---|---|---|---|---|---|
| C01 | 21 | 59.24–62.19 | 58.99–62.44 | 4 (ZFST+XP-EHH+Rsb+π) | Select; π↓ Select. SERPINA1, SERPINA3-1, SERPINA3-3, SERPINA5, SERPINA6, SERPINA11 … 106 QTL (Health; Milk; Exterior) |
| C02 | 1 | 85.79–86.15 | 85.54–86.40 | 3 (XP-EHH+Rsb+π) | Control; π↓ Control. CCDC39, DNAJC19, FXR1, TTC14. 17 QTL (Milk; Production; Reproduction) |
| C03 | 3 | 26.06–26.63 | 25.81–26.88 | 3 (XP-EHH+Rsb+π) | Control; π↓ Control. CD2, CD58, CD101, IGSF3, MAN1A2, PTGFRN … 39 QTL (Milk; Production; Reproduction) |
| C04 | 3 | 31.37–32.19 | 31.12–32.44 | 3 (ZFST+XP-EHH+π) | Control; π↓ Control. KCND3, RAP1A, CHIA, OVGP1, ATP5PB, CEPT1 … 31 QTL (Reproduction; Carcass; Production) |
| C05 | 3 | 46.37–48.33 | 46.12–48.58 | 3 (ZFST+Rsb+π) | Select; π↓ Control. DPYD, ALG14, CNN3, PTBP2, RWDD3, TLCD4. 72 QTL (Milk; Production; Reproduction) |
| C06 | 5 | 2.25–3.04 | 2.00–3.29 | 3 (XP-EHH+Rsb+π) | Select; π↓ Select. TRHDE. 40 QTL (Milk; Production; Health) |
| C07 | 6 | 82.58–82.77 | 82.33–83.02 | 3 (XP-EHH+Rsb+π) | Select; π↓ Select. no named genes. 268 QTL (Milk; Health; Carcass) |
| C08 | 7 | 57.70–58.86 | 57.45–59.11 | 3 (XP-EHH+Rsb+π) | Select; π↓ Select. SPINK1, DPYSL3, GPR151, JAKMIP2, POU4F3, PPP2R2B … 39 QTL (Milk; Carcass; Production) |
| C09 | 8 | 29.88–30.52 | 29.63–30.77 | 3 (XP-EHH+Rsb+π) | Select; π↓ Control. NFIB. 21 QTL (Milk; Exterior; Health) |
| C10 | 9 | 96.69–97.80 | 96.44–98.05 | 3 (ZFST+Rsb+π) | Select; π↓ Select. PLG, SLC22A3, AGPAT4, MAP3K4, PRKN. 47 QTL (Milk; Carcass; Reproduction) |
| C11 | 12 | 71.32–73.10 | 71.07–73.35 | 3 (XP-EHH+Rsb+π) | Select; π↓ Select. CLDN10, DNAJC3, DZIP1, UGGT2. 39 QTL (Milk; Exterior; Health) |
| C12 | 15 | 24.76–25.37 | 24.51–25.62 | 3 (XP-EHH+Rsb+π) | Select; π↓ Select. ZBTB16, C15H11orf71, HTR3A, NXPE2, NXPE4, RBM7 … 29 QTL (Carcass; Production; Milk) |
| C13 | 15 | 27.51–28.84 | 27.26–29.09 | 3 (ZFST+Rsb+π) | Select; π↓ Select. APOA1, APOA4, APOA5, APOC3, CD3D, CD3E … 154 QTL (Milk; Health; Exterior) |
| C14 | 16 | 47.78–48.79 | 47.53–49.04 | 3 (XP-EHH+Rsb+π) | Select; π↓ Select. AJAP1. 22 QTL (Milk; Health; Carcass) |
| C15 | 16 | 77.61–78.83 | 77.36–79.08 | 3 (XP-EHH+Rsb+π) | Select; π↓ Control. PTPRC, NR5A2, ATP6V1G3. 146 QTL (Health; Carcass; Milk) |
| C16 | 23 | 2.21–2.40 | 1.96–2.65 | 3 (XP-EHH+Rsb+π) | Control; π↓ Control. PRIM2. 7 QTL (Milk; Production; Exterior) |
| C17 | 26 | 27.68–27.72 | 27.43–27.97 | 3 (XP-EHH+Rsb+π) | Select; π↓ Select. SORCS1. 314 QTL (Milk; Health; Carcass) |

**Table 3.** Sensitivity of the line difference in autozygosity to ROH-calling parameters.

| **Caller** | **Min markers** | **Min length** | **Control F_ROH** | **Select F_ROH** | **Excess** | **P** |
|---|---|---|---|---|---|---|
| detectRUNS | 20 | 250 kb | 0.1171 | 0.1318 | +0.0147 (+12.5%) | 1.3 × 10<sup>−18</sup> |
| PLINK v1.9 | 30 | 500 kb | 0.0784 | 0.0942 | +0.0158 (+20.2%) | 7.9 × 10<sup>−21</sup> |
| PLINK v1.9 | 50 | 1,000 kb | 0.0625 | 0.0779 | +0.0154 (+24.7%) | 9.4 × 10<sup>−22</sup> |
| PLINK v1.9 | 75 | 2,000 kb | 0.0531 | 0.0678 | +0.0147 (+27.7%) | 2.5 × 10<sup>−22</sup> |

*The absolute excess is stable across an eight-fold range of minimum length and two callers; the relative excess grows as the threshold rises, because the difference lies in long ROH.*

## Discussion

Unlike studies that rely on breed contrasts or pseudo-controls, the Langhill design lets us attribute genomic signatures directly to sire selection; their convergence across complementary statistics, and their coincidence with the herd’s recorded fertility and health responses, reinforces that attribution. The Langhill experiment demonstrates, at the level of the genome, the dual character of intensive dairy selection. Five decades of sire selection produced clear phenotypic gains accompanied by a coherent set of genomic signatures: slower LD decay, elevated ROH, a modest genome-wide reduction in nucleotide diversity, and a discrete set of differentiated regions. Together these describe a Select genome that is more structured and less diverse than its Control counterpart.

Earlier comparisons of selected and unselected Holstein lines, notably the University of Minnesota lines established in 1964^15,16^, contrasted selection against no selection. Langhill instead contrasts elite with average sires drawn from the same contemporaneous national population, which is the choice breeders actually face. Within that design, several results are, to our knowledge, new. The Select line’s excess autozygosity is confined to long, recent ROH and is not concentrated at any locus. It arose although the Select line used more sires, and is explained by higher co-ancestry among elite sires rather than by intensive use of a few bulls. It was already present in the earliest genotyped cohort and thereafter accumulated at the same annual rate as in the Control line, while nucleotide diversity declined across cohorts only in the Select line. Four of the seventeen convergent regions carried the haplotype and diversity signal in the Control line, so a line sired by average-merit bulls is not a static reference: it too changes, through drift or selection, and contrasts with it measure divergence between two moving lines. Finally, because Control-line daughters were sired by markedly older bulls, the two lines differ in generation interval, so per-generation estimates of Ne differ between lines even though annual rates of inbreeding do not, a caution for comparing Ne across selection lines or breeding programmes.

### Erosion of genetic diversity under directional selection

Over five decades, continuous directional sire selection in this experimental herd has driven substantial phenotypic and genetic gains, but these have been accompanied by a clear and measurable erosion of genetic diversity. We quantified this cost through a coherent set of genomic signatures: linkage disequilibrium (LD) extends further across all measured distances, overall genomic relatedness is threefold higher, and the Select line exhibits a 24.7% excess in autozygosity. These changes accumulated in a closed herd with an effective population size (*N_e_*) of ∼33. This is low, but of the same order as estimates for commercial Holstein populations, which commonly lie around or below 100^24,25^. The annual rate of inbreeding does not distinguish the lines, and the higher per-generation estimate for the Select line (43 versus 26) reflects its shorter generation interval rather than greater diversity. What distinguishes the lines is the level of autozygosity. Crucially, the length-resolved decomposition of runs of homozygosity (ROH) allows us to precisely date this excess. Because ROH shorter than 2 Mb do not differ between the Select and Control lines, while every longer class differs significantly, we can conclude that the additional autozygosity is entirely recent. It was accrued in the generations following the divergence of the lines rather than inherited from a shared ancestral base population; the offset was already established in the earliest genotyped cohort (born 1990) and has since been maintained rather than enlarged.

### Linking selection signatures to the traits under selection

When the genomic evidence is examined locus by locus, the different methods divide sharply. Frequency- and haplotype-based selection scans successfully converge on a discrete, limited set of genomic regions, most notably on BTA21 (59.2–62.2 Mb; four methods) and BTA3 (31.4–32.2 Mb; three methods), with 15 further three-method regions (Table 2). Because the traits under selection are highly polygenic, these regions most likely account for only a small proportion of trait variance: the scans do not resolve the architecture of these traits, but link the line contrast to regions already associated with them. Where selection has raised the frequency of favourable alleles in these regions, the same signatures mark the genomic benefit of selection, the counterpart to the loss of diversity described above. Autozygosity, conversely, carries no locus-specific information in this population; it is elevated genome-wide, yet no single locus exceeds the background threshold. Rather than being contradictory, these findings are highly complementary, capturing two fundamentally distinct evolutionary processes. Intense directional selection has left localized footprints where it acted directly upon specific, advantageous haplotypes. Inbreeding, however, accumulated diffusely across the genome as a demographic by-product of the selection regime. Consequently, ROH serve as an index of the latter demographic process and, in the context of this herd, should not be conflated with a map of selection targets.

### The Demographic Mechanism Driving Autozygosity

The origin of this excess autozygosity presents a highly counterintuitive demographic scenario. The Select line was actually sired by *more* individual bulls than the Control line (79 versus 72 sires for 543 and 583 genotyped cows, respectively). Furthermore, each sire in the Select line contributed fewer daughters on average (6.9 vs 8.1), the top five most heavily used sires accounted for a smaller total share of daughters (16.8% vs 18.9%), and the line experienced fewer close matings (1.46% vs 1.61% of pairs at a kinship > 0.125).

What mechanistically distinguishes the Select line is that its cows are intrinsically three times more related to one another. Because elite UK Holstein sires trace their lineage back to an exceptionally narrow set of ancestors, selecting strictly on top-decile genetic merit invariably samples from a genealogically restricted pool—regardless of how many individual bulls are incorporated into the breeding program. This evidence demonstrates that the autozygosity cost of elite sire selection is incurred primarily through the shared co-ancestry of the elite pool, rather than through the intensive use of a few individual bulls. This distinction carries direct, practical consequences for the optimization of modern mating designs, highlighting the limitation of managing inbreeding solely by restricting individual sire contributions.

The convergent regions are the most credible selection candidates because they are recovered by several complementary statistics (which are computed from the same genotypes and so are not statistically independent). Annotation of the 17 regions supported by three or more scans (Table 2; Tables S8–S9) identified several biologically coherent intervals. The four-method region on BTA21 (59.2–62.2 Mb) spans the SERPINA cluster of serine-protease inhibitors (SERPINA1, SERPINA3 paralogues, SERPINA5, SERPINA6 and others), with 106 curated QTL chiefly for health, milk and conformation traits. Select-line three-method regions include the BTA15 interval at 27.5–28.8 Mb carrying the apolipoprotein cluster (APOA1, APOA4, APOA5, APOC3) and the CD3 T-cell receptor genes (CD3D, CD3E, CD3G) with IL10RA (154 QTL, led by somatic cell score, milk protein percentage and milk fatty-acid traits); a BTA6 interval at 82.6–82.8 Mb, about 2.6 Mb proximal to the casein cluster (CSN1S1 at 85.4 Mb), carrying 268 QTL, led by milk κ-casein percentage; PTPRC (CD45) on BTA16; and SORCS1 on BTA26 (314 QTL for milk fatty-acid indices). Immune-related genes thus recur among the convergent regions. Because a single study can contribute thousands of QTLdb entries to one interval (here 9,439 entries from one study in a single BTA6 region), we tested enrichment counting each trait–publication pair once (Table S10): health QTL were modestly over-represented among candidate regions (about 1.3-fold; FDR < 0.01), whereas milk and production QTL were not, and no individual trait remained enriched after correction. None of the classical large-effect milk genes — DGAT1^26^, ABCG2^27^, the casein cluster or GHR — lies within any candidate region, consistent with these variants having been at high frequency in elite and average bulls alike before the lines diverged. Gene-level causality cannot be inferred from array-resolution intervals. The overlap of selection signals with fertility loci echoes the U.S. Holstein finding that milk-yield selection dragged on fertility and immunity through hitchhiking^15^, consistent with the documented fertility decline in the Langhill Select line^12,28^.

For the sustainability of dairy farming, the reduction in diversity and rise in homozygosity in the Select line is the central message. Reduced diversity can constrain future response to selection and may increase susceptibility to disease and environmental stress^13,29,30^. The Control line is therefore valuable as a reservoir of variation and a benchmark for diversity, and the experiment argues for breeding strategies that actively manage diversity (e.g. optimal-contribution selection) alongside genetic gain.

## Limitations

Genotypes derive from an ∼80K array imputed across platforms; the sparse marker set limits fine-scale resolution and makes single-window π-ratios unreliable, so haplotype-based and cross-method signals carry more weight. Langhill is a small, closed herd with declining size, inflating drift relative to selection; some regions may reflect drift, which the cross-method intersection mitigates but cannot fully resolve. The ZFST, XP-EHH and Rsb scans used the 1,073 cows whose recorded line agreed with genomic clustering; this raises mean FST (0.023 versus 0.020 on all 1,126 cows) but leaves the differentiated regions essentially unchanged. The XP-EHH and Rsb scans depend on haplotype phase: the cows were phased separately within line, and re-phasing the same genotypes jointly across both lines produced no haplotype signal, so these two scans are treated as supporting rather than stand-alone evidence. Counts of markers beyond |4| are not in themselves a test of selection, because random splits of a single line also yield many such markers; we therefore relied on replication of the signal across random subsamples of the cows and on its coincidence with the ZFST and π scans (Supplementary Note S1). Most extreme haplotype scores were higher in the Select line, as expected from its higher autozygosity and relatedness. The BTA21 and BTA3 intervals are supported by ZFST and π independently of phase; the BTA10 22.3 Mb XP-EHH peak was less stable and should be regarded as provisional.

Finally, while the design involves potential confounders such as herd relocation and unmeasured environmental stressors, the study’s validity remains robust. Feeding systems were balanced across lines among genotyped cows (Control: 270/274; Select: 250/280; *χ*^2^ = 0.56, *P* = 0.46), ensuring the line contrast is not carried by diet. Furthermore, while imputed genotypes may inflate absolute *F_ROH_* values, this affects the overall level rather than the comparative contrast. Given that poorer imputation quality typically increases apparent homozygosity, the fact that the less-represented Control line exhibits *less* autozygosity confirms that our results remain conservative and are not artifacts of the imputation process.

### Breeding implications

Selection decisions in modern dairy programmes are, and will remain, driven by the estimated breeding values of individual animals; these results do not argue otherwise. Their relevance is to mating decisions, where diversity is actually managed. That the Select line accumulated a quarter more autozygosity than its contemporaneous control while using more sires shows that counting bulls is an inadequate safeguard: an elite pool can be genealogically narrow however broadly it is sampled. Optimal-contribution selection and mate-allocation schemes that constrain the rate of co-ancestry rather than the number of sires target the mechanism operating here^31,32^. That the excess is confined to long, recent ROH is encouraging in one respect: it is the component most amenable to management through mating decisions on a generation-by-generation basis, unlike founder autozygosity, which is already fixed.

### Future directions

Feeding-system design in the studied population suits dissection of genotype × environment × management (G×E×M) interactions. Priorities are: functional follow-up of the convergent regions, beginning with the BTA21 SERPINA cluster, the BTA15 apolipoprotein and CD3 cluster and the chr9 ∼97.7 Mb locus; whole-genome resequencing of archived samples to overcome array resolution; modelling the temporal trajectory of diversity and inbreeding; integration with fertility and health records to test whether differentiated regions co-locate with fitness QTL; and extension of all analyses to the approximately 2,300 Langhill cows now genotyped.

## Conclusion

The Langhill experiment demonstrates, genomically, that sustained selection for yield delivers real phenotypic gains but at a measurable cost to diversity and homozygosity. Long-term experiments of this kind are an irreplaceable resource for quantifying that trade-off and for designing breeding programmes that are both productive and resilient.

## Methods

### Animals and selection lines

The Langhill Holstein-Friesian herd (established in 1973; SRUC Dairy Research Centre, Crichton Royal Farm) comprises a Select line of cows, sired by bulls in the top decile of UK genetic merit for milk fat-plus-protein yield, and a Control line, sired by bulls of UK-average merit, with a nested high/low feeding contrast from 1988. Sires were ranked on predicted transmitting ability (PTA) for kg fat plus protein, and matings to close relatives were avoided. Bulls used in the Select line are not subsequently used in the Control line, even as they age and decline in relative merit. Line membership was taken from the herd breeding records; no genomic criterion was used to assign line, and genomic clustering was used to exclude animals only from the ZFST, XP-EHH and Rsb scans (see Genetic differentiation). A total of 1,126 cows with a recorded line (583 Control, 543 Select) were included in the present study. Cows were born between 1990 and 2016; numbers of cows, sires and dams per line and birth-year band are given in Table 4. Cows were milked three times daily; morning, midday and evening yields sum to total daily yield. Herd records document occasional historical parentage errors arising from mis-assigned insemination straws; affected animals are excluded from the trial when identified. Because such errors mix the two gene pools, any residual cases attenuate rather than inflate between-line contrasts, so the differences reported here are conservative.

**Table 4.** Genotyped cows, sires and dams per line and birth-year band.

| Birth years | Control cows | Select cows | Control sires | Select sires | Control dams | Select dams |
|---|---|---|---|---|---|---|
| 1990–1994 | 17 | 28 | 10 | 12 | 17 | 27 |
| 1995–1999 | 81 | 44 | 15 | 10 | 65 | 40 |
| 2000–2004 | 127 | 88 | 18 | 18 | 96 | 72 |
| 2005–2009 | 214 | 158 | 20 | 15 | 158 | 122 |
| 2010–2016 | 144 | 225 | 23 | 30 | 113 | 159 |
| Total | 583 | 543 | 72 | 79 | 396 | 380 |
| Daughters per sire | 8.1 | 6.9 | — | — | — | — |
| Top-5 sires, % of daughters | 18.9% | 16.8% | — | — | — | — |

*The Select line used more sires, each with fewer daughters. Sires and dams may appear in more than one band, so those columns sum to more than the line total.*

**Table 5.** Autozygosity by ROH length class (ROH called at 30 SNPs / 500 kb).

| **Length class** | **Control F_ROH** | **Select F_ROH** | **Excess** | **P** | **% of total excess** |
|---|---|---|---|---|---|
| 0.5–1 Mb | 0.0027 | 0.0027 | +2.3% | 0.29 (n.s.) | 0.4% |
| 1–2 Mb | 0.0139 | 0.0140 | +0.8% | 0.45 (n.s.) | 0.7% |
| 2–4 Mb | 0.0148 | 0.0176 | +18.5% | 1.7 × 10<sup>−15</sup> | 17.3% |
| 4–8 Mb | 0.0168 | 0.0210 | +25.1% | 8.4 × 10<sup>−19</sup> | 26.6% |
| 8–16 Mb | 0.0171 | 0.0220 | +28.5% | 4.2 × 10<sup>−14</sup> | 30.8% |
| >16 Mb | 0.0131 | 0.0169 | +29.3% | 2.1 × 10<sup>−5</sup> | 24.1% |
| Total | 0.0784 | 0.0942 | +20.2% | 7.9 × 10<sup>−21</sup> | 100% |

*Short ROH (founder ancestry) do not differ between lines; the excess is confined to long, recent ROH.*

### Phenotypic comparisons

Phenotypes were a single test-day record per cow for the 1,348 cows with a recorded line (675 Control, 673 Select), including 222 cows that were not genotyped. Lines were compared for total daily milk yield, its morning, midday and evening components, daily protein and fat yield, body weight, body condition score and stage of lactation (week of lactation at the test day) by two-sided Mann–Whitney tests. Zero values in the yield traits were treated as missing, following the herd recording convention: protein and fat yield carry 413 such zeros (31% of records), and their inclusion distorts both means and tests.

### Genotyping and imputation

DNA extracted from archived whole blood (DNeasy Blood & Tissue kit, QIAGEN) had been genotyped previously^33^ on the Illumina BovineSNP50, GGP Bovine 50K and GGP Bovine HD 150K and imputed to the 80K UK national-evaluation panel with FindHap 3^34^. The analysed file contains 76,991 markers on 1,227 animals with a post-imputation call rate of 1.00. Autosomes 1–29 were analysed (74,375 markers; ARS-UCD1.2), spanning 2,479,364 kb of marker-covered sequence at a mean spacing of 33.3 kb. Markers coded 30 and 31 — labelled X and Y under PLINK’s --chr-set 29 convention — were excluded: the two groups are contiguous (ending 133,289,707 bp and beginning 133,312,403 bp) and together span 138.9 Mb, corresponding to the bovine X chromosome (ARS-UCD1.2: 139.0 Mb) divided at the pseudoautosomal boundary. ARS-UCD1.2 contains no Y assembly, so no Y-linked markers are present. Two samples (G102, G384; Table S7) were flagged by genomic relatedness checks: neither shared kinship with its recorded paternal half-sibs (KING kinship < 0; expected ≈ 0.125) while both had first-degree relatives in the opposite line, indicating possible sample misidentification. Both are retained among the 1,126 cows analysed; excluding them altered no result (for example, the detectRUNS F_ROH excess was +0.0147 with or without them). Genotypes are recorded as A/B alleles recoded to A/T, so strand and reference-allele checks are not possible; this does not affect dosage-based analysis.

### Population structure and linkage disequilibrium

Markers were LD-pruned with PLINK 2.0^35^ (--indep-pairwise 50 5 0.2), retaining 17,852. Principal components were computed on the pruned set (PLINK 2.0 --pca, --nonfounders) and ancestry proportions with fastSTRUCTURE^36^ (K = 1–10; K = 2 shown), animals assigned to a cluster at Q > 0.6. Genomic relatedness was estimated as KING-robust kinship^37^ on the same pruned set. LD decay (r²) was computed within 300 kb per line and summarised in 2-kb bins weighted by pair count, with 95% confidence intervals from the pair-weighted between-distance variance within each bin.

### Nucleotide diversity

Nucleotide diversity (π)^38^ was computed per line in 10-kb windows (VCFtools^39^) and additionally stratified by birth cohort. Because 93% of windows contain a single marker, adjacent windows are not independent; cohort means are therefore compared by regression on cohort (n = 5 per line) and confidence intervals obtained by resampling chromosomes with replacement (2,000 replicates). Window-level rank tests are not reported. The ratio log₂(π_Select/π_Control) of 25-window centred rolling means of π (all 1,126 cows) was scanned in the 0.5% tails.

### Autozygosity

Runs of homozygosity^40^ were called in PLINK v1.9^41^ on unpruned autosomal genotypes — LD pruning was not applied, as it destroys the marker contiguity on which ROH detection depends — with: --homozyg-snp 50, --homozyg-kb 1000, --homozyg-density 100, --homozyg-gap 1000, --homozyg-window-snp 50, --homozyg-window-het 1, --homozyg-window-missing 5, --homozyg-window-threshold 0.05, --homozyg-het 1. The minimum marker count was derived rather than assumed: with mean expected heterozygosity 0.4013 across 74,375 markers in 1,126 animals, the threshold keeping the expected number of false-positive runs below 0.05 per genome is l = log(α/(n_s·n_i))/log(1−het) = 41.4, i.e. 42 markers; 50 was used. F_ROH is total autosomal ROH length divided by the marker-covered autosomal length (2,479,364 kb); cows with no detected ROH would contribute F_ROH = 0 (none occurred). Sensitivity to parameter choice was assessed by re-calling at 30 markers/500 kb and 75 markers/2,000 kb, and against detectRUNS sliding-window calling^42^ (15-SNP windows, at most one heterozygous and one missing genotype per window, 20 markers/250 kb; Table 3), applied to the same 1,126 cows, autosomes and denominator. ROH were assigned to length classes (0.5–1, 1–2, 2–4, 4–8, 8–16, >16 Mb) and lines compared per class by Mann–Whitney test. To test whether autozygosity was concentrated at any locus, each cow was scored as autozygous at a marker if it fell within a ROH of at least 2 Mb and the Select-versus-Control odds ratio computed per marker; because genome-wide F_ROH differs between lines, odds ratios were compared not against unity but against an empirical null generated by permuting line within deciles of each cow’s genome-wide long-ROH burden (200 permutations), preserving the genome-wide difference while breaking any locus-specific association.

Effective population size was estimated from the rate of inbreeding^43^. Individual F_ROH was regressed on birth year separately by line and with a year × line interaction; because the interaction was not significant, a common slope was fitted. The slope was converted to a per-generation rate using the mean generation interval measured from the pedigree, 6.8 years, and Ne obtained as 1/(2ΔF). The generation interval is the mean of the dam pathway (3.77 years; 1,109 dam–daughter pairs) and the sire pathway (9.86 years; 961 sire–daughter pairs, using birth dates available for 118 of the 146 sires). By line it was 8.2 years in the Control line (sire pathway 12.5 years) and 5.4 years in the Select line (sire pathway 7.2 years), reflecting the use of older, progeny-proven bulls in the Control line. Summarising the pathways by medians gives 5.9 years (Ne ≈ 38), and the 5.4-year interval typical of dairy populations gives Ne ≈ 42; these are reported as sensitivity estimates. An LD-based estimator was not used because the generations it samples scale as t = 1/(2c), so the available 300-kb window reaches no closer than ∼167 generations — an order of magnitude beyond the duration of the experiment.

### Genetic differentiation

The ZFST, XP-EHH and Rsb scans were run on the 1,073 cows whose recorded line agreed with the K = 2 clustering (577 Control, 496 Select), excluding the 53 discordant cows; the π scan used all 1,126 cows. Because 41 of the 53 were Select cows born before 2000, this exclusion sharpens the line contrast; the ZFST scan was therefore repeated on all 1,126 cows. Per-SNP FST (Weir & Cockerham^44^) was computed between lines in VCFtools^39^, smoothed as a centred 10-SNP rolling mean and Z-standardised (ZFST); markers with ZFST > 5 were merged into regions within 1 Mb.

### Haplotype-based scans and cross-method convergence

XP-EHH and Rsb (Select versus Control) were computed with rehh^21^, with |score| > 4 defining outlier markers. Haplotypes were phased separately within each line (496 Select and 577 Control cows); rehh was run with default settings (minor-allele-frequency threshold 0.05) and scores were standardised genome-wide. Because these statistics depend on the phase, they are interpreted alongside the allele-frequency and diversity scans (Limitations; Supplementary Note S1). Regions were formed within each scan by merging outlier markers (ZFST, XP-EHH, Rsb) or tail windows of the π ratio (Control > Select and Select > Control) lying within 1 Mb, and regions from different scans within 1 Mb of one another were combined into 132 candidate regions, each scored for the scans supporting it (Figure 5; Table S6). Candidate regions with ±250 kb flanks were annotated with GALLO^23^ for Ensembl release 104 genes on ARS-UCD1.2 and for Animal QTLdb^22^ QTL (cattle release 59, whose positions match the ARS-UCD1.2 release-76 file for 99.97% of 160,463 shared QTL). Because single studies can contribute thousands of QTLdb entries to one interval, QTL enrichment was tested counting each trait–publication pair once, by one-sided hypergeometric tests against all autosomal QTLdb entries with Benjamini–Hochberg correction (Table S10).

## Supplementary information

**Figure S1.**
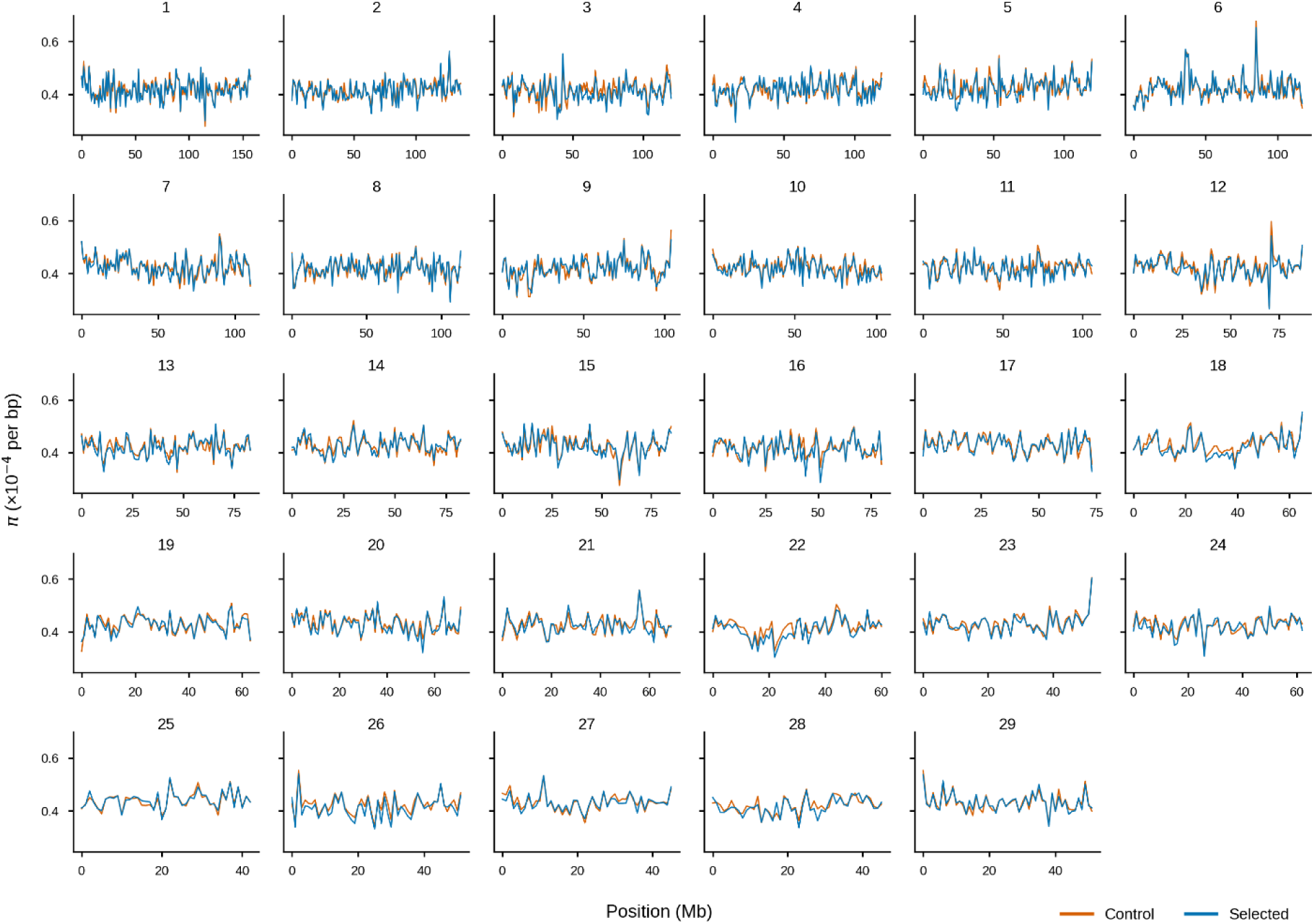
Nucleotide diversity along each autosome. Nucleotide diversity (π, ×10⁻⁴ per bp) against position (Mb) for the Control and Select lines, autosomes 1–29.

Supplementary tables: S1 per-cohort π by line with block-bootstrap confidence intervals; S2 segment count and mean segment length per cow by line; S3 LD decay with 95% CI by 2-kb bin and line; S4 full ZFST region list (28); S5 full XP-EHH (32) and Rsb (73) region lists; S6 cross-method intersection matrix across 132 candidate regions; S7 flagged samples (G102, G384) with KING kinship to recorded half-sibs and to first-degree relatives in the opposite line; S8 genes within ±250 kb of each candidate region (GALLO, Ensembl 104); S9 Animal QTLdb QTL within ±250 kb of each candidate region; S10 publication-aware QTL enrichment. Supplementary Note S1 (separate file) describes the robustness checks on the XP-EHH and Rsb scans.

## List of abbreviations

BCS: body condition score
F_ROH: proportion of the autosomal genome in runs of homozygosity
F_ST: fixation index
LD: linkage disequilibrium
PC: principal component
QTL: quantitative trait locus
ROH: runs of homozygosity
Rsb: ratio of extended haplotype homozygosity between populations
SNP: single nucleotide polymorphism
XP-EHH: cross-population extended haplotype homozygosity
ZF_ST: Z-transformed F_ST
π: nucleotide diversity.

## Declarations

### Ethics approval and consent to participate

Blood samples were collected from the Langhill herd at the SRUC Dairy Research Centre in accordance with UK Home Office regulations (PPL 60/4278) and with the approval of the SRUC Animal Experimentation Committee, as reported for the same archived samples by Ilska-Warner et al.^33^ The present study used these archived samples together with routine herd records. The Langhill herd is managed under normal husbandry, with individual feed intake recorded, and the phenotype and pedigree data analysed here are routine herd records; no additional procedures were carried out on animals for this study.

### Consent for publication

Not applicable.

### Availability of data and materials

The data and code needed to reproduce Figures 1–5 are deposited in Zenodo (DOI 10.5281/zenodo.24005571). The deposit consists of one Excel workbook (Langhill_data.xlsx) and one R script (make_figures.R). The workbook holds the anonymised per-animal data (line, birth year, feeding group, F_ROH and ROH length classes, principal components and ancestry coefficients), with animals identified only by anonymous codes (G001, G002,…), together with the milk-yield records and the per-marker, per-window and per-region results from which every figure is drawn; the script redraws Figures 1–5 and recomputes the statistics quoted in their legends. Supplementary Tables S1–S10 and Supplementary Note S1 accompany this article. Analyses used PLINK v1.9 and v2.0, VCFtools, rehh, fastSTRUCTURE and GALLO; coordinates are ARS-UCD1.2 throughout. The individual-animal genotype data from the Langhill herd cannot be made publicly available and are not part of the deposit; they are held by SRUC, and requests for access can be made to the corresponding author and will be considered under SRUC data-sharing procedures.

### Use of AI tools

Claude (Anthropic) was used to help write and check analysis scripts, to cross-check numbers against the source files and to edit the text. The authors reviewed all output, ran the analyses and take full responsibility for the content.

### Authors’ contributions

All authors contributed to the conception and design of the study, to the analysis and interpretation of the results and to the critical revision of the manuscript. RS performed the analyses and drafted the manuscript. All authors read and approved the final manuscript.

## Acknowledgements

We acknowledge the SRUC Dairy Research Centre (Crichton Royal Farm) for access to the Langhill herd records and archived samples.

## References

1. Mehrabi, Z. et al. Livestock policy for sustainable development. Nature Food 1, 160–165 (2020). 10.1038/s43016-020-0042-9

2. Van Zanten, H.H.E., Van Ittersum, M.K. & De Boer, I.J.M. The role of farm animals in a circular food system. Global Food Security 21, 18–22 (2019). 10.1016/j.gfs.2019.06.003

3. Thompson, P.B. & Nardone, A. Sustainable livestock production: methods and ethics. Livestock Production Science 61, 111–119 (1999). 10.1016/S0301-6226(99)00062-7

4. Bilotto, F. et al. Towards resilient, inclusive, sustainable livestock farming systems. Trends in Food Science & Technology 152, 104668 (2024). 10.1016/j.tifs.2024.104668

5. Mottet, A. et al. Livestock: on our plates or eating at our table? Global Food Security 14, 1–8 (2017). 10.1016/j.gfs.2017.01.001

6. Opio, C. et al. Greenhouse Gas Emissions from Ruminant Supply Chains – A Global Life Cycle Assessment. FAO, Rome (2013). ISBN 978-92-5-107945-4.

7. Havlík, P. et al. Climate change mitigation through livestock system transitions. PNAS 111, 3709–3714 (2014). 10.1073/pnas.1308044111

8. Bajželj, B. et al. Importance of food-demand management for climate mitigation. Nature Climate Change 4, 924–929 (2014). 10.1038/nclimate2353

9. Palma-Vera, S.E. et al. Genomic characterization of the world’s longest selection experiment in mouse reveals the complexity of polygenic traits. BMC Biology 20, 52 (2022). 10.1186/s12915-022-01248-9

10. Moose, S.P., Dudley, J.W. & Rocheford, T.R. Maize selection passes the century mark: a unique resource for 21st century genomics. Trends in Plant Science 9, 358–364 (2004). 10.1016/j.tplants.2004.05.005

11. Hill, W.G. Can more be learned from selection experiments of value in animal breeding programmes? Or is it time for an obituary. J. Anim. Breed. Genet. 128, 87–94 (2011). 10.1111/j.1439-0388.2010.00913.x

12. Pryce, J.E., Nielsen, B.L., Veerkamp, R.F. & Simm, G. Genotype and feeding system effects and interactions for health and fertility traits in dairy cattle. Livestock Production Science 57, 193–201 (1999). 10.1016/S0301-6226(98)00180-8

13. Coffey, M.P., Simm, G. & Brotherstone, S. Energy balance profiles for the first three lactations of dairy cows estimated using random regression. J. Dairy Sci. 85, 2669–2678 (2002). 10.3168/jds.S0022-0302(02)74352-X

14. Liu, X. et al. Selection signature analysis of whole-genome sequences to identify genome differences between selected and unselected Holstein cattle. Animals 15, 2247 (2025). 10.3390/ani15152247

15. Ma, L. et al. Genome changes due to artificial selection in U.S. Holstein cattle. BMC Genomics 20, 128 (2019). 10.1186/s12864-019-5459-x

16. Kim, E.-S., Cole, J.B., Huson, H. et al. Effect of artificial selection on runs of homozygosity in U.S. Holstein cattle. PLoS ONE 8, e80813 (2013). 10.1371/journal.pone.0080813

17. Bertolini, F. et al. Exploring genome-wide differentiation and signatures of selection in Italian and North American Holstein populations. J. Dairy Sci. 106, 5829–5847 (2023). 10.3168/jds.2022-22941

18. Korkuć, P. et al. Genomic diversity and selection signatures of the German Black Pied (DSN) breed relative to Holstein. Frontiers in Genetics 13, 993959 (2022). 10.3389/fgene.2022.993959

19. Zhang, Q. et al. Identification of selective sweeps in the Chinese Holstein and Simmental. BMC Genomics 16, 542 (2015). 10.1186/s12864-015-1715-x

20. Wall, E., Brotherstone, S., Woolliams, J.A., Banos, G. & Coffey, M.P. Genetic evaluation of fertility using direct and correlated traits. J. Dairy Sci. 86, 4093–4102 (2003). 10.3168/jds.S0022-0302(03)74023-5

21. Gautier, M., Klassmann, A. & Vitalis, R. rehh 2.0: a reimplementation of the R package rehh to detect positive selection from haplotype structure. Mol. Ecol. Resour. 17, 78–90 (2017). 10.1111/1755-0998.12634

22. Hu, Z.-L., Park, C.A. & Reecy, J.M. Bringing the Animal QTLdb and CorrDB into the future. Nucleic Acids Res. 50, D956–D961 (2022). 10.1093/nar/gkab1116

23. Fonseca, P.A.S. et al. GALLO: an R package for genomic annotation and integration of multiple data sources in livestock for positional candidate loci. GigaScience 9, giaa149 (2020). 10.1093/gigascience/giaa149

24. Rodríguez-Ramilo, S.T., Fernández, J., Toro, M.A., Hernández, D. & Villanueva, B. Genome-wide estimates of coancestry, inbreeding and effective population size in the Spanish Holstein population. PLoS ONE 10, e0124157 (2015). 10.1371/journal.pone.0124157

25. Salehi, R., Javanmard, A., Mokhber, M. & Alijani, S. Estimating linkage disequilibrium and effective population size across generations in Holstein cattle. Vet. Med. Sci. 11, e70684 (2025). 10.1002/vms3.70684

26. Grisart, B. et al. Positional candidate cloning of a QTL in dairy cattle: identification of a missense mutation in the bovine DGAT1 gene with major effect on milk yield and composition. Genome Res. 12, 222–231 (2002). 10.1101/gr.224202

27. Cohen-Zinder, M. et al. Identification of a missense mutation in the bovine ABCG2 gene with a major effect on the QTL on chromosome 6 affecting milk yield and composition in Holstein cattle. Genome Res. 15, 936–944 (2005). 10.1101/gr.3806705

28. Banos, G. & Coffey, M.P. Genetic association between body energy measured throughout lactation and fertility in dairy cattle. Animal 4, 189–199 (2010). 10.1017/S1751731109991182

29. Notter, D.R. The importance of genetic diversity in livestock populations of the future. J. Anim. Sci. 77, 61–69 (1999). 10.2527/1999.77161x

30. Banos, G., Coffey, M.P., Wall, E. & Brotherstone, S. Genetic relationship between first-lactation body energy and later-life udder health in dairy cattle. J. Dairy Sci. 89, 2222–2232 (2006). 10.3168/jds.S0022-0302(06)72293-7

31. Sonesson, A.K. & Meuwissen, T.H.E. Mating schemes for optimum contribution selection with constrained rates of inbreeding. Genet. Sel. Evol. 32, 231–248 (2000). 10.1186/1297-9686-32-3-231

32. Meuwissen, T.H.E. Maximizing the response of selection with a predefined rate of inbreeding. J. Anim. Sci. 75, 934–940 (1997). 10.2527/1997.754934x

33. Ilska-Warner, J.J., Psifidi, A., Seeker, L.A., Wilbourn, R.V., Underwood, S.L., Fairlie, J., Whitelaw, B., Nussey, D.H., Coffey, M.P. & Banos, G. The genetic architecture of bovine telomere length in early life and association with animal fitness. Frontiers in Genetics 10, 1048 (2019). 10.3389/fgene.2019.01048

34. VanRaden, P.M., Null, D.J., Sargolzaei, M., Wiggans, G.R., Tooker, M.E., Cole, J.B. et al. Genomic imputation and evaluation using high-density Holstein genotypes (FindHap). J. Dairy Sci. 96, 668–678 (2013). 10.3168/jds.2012-5702

35. Chang, C.C. et al. Second-generation PLINK: rising to the challenge of larger and richer datasets. GigaScience 4, 7 (2015). 10.1186/s13742-015-0047-8

36. Raj, A., Stephens, M. & Pritchard, J.K. fastSTRUCTURE: variational inference of population structure in large SNP datasets. Genetics 197, 573–589 (2014). 10.1534/genetics.114.164350

37. Manichaikul, A. et al. Robust relationship inference in genome-wide association studies. Bioinformatics 26, 2867–2873 (2010). 10.1093/bioinformatics/btq559

38. Nei, M. & Li, W.-H. Mathematical model for studying genetic variation in terms of restriction endonucleases. PNAS 76, 5269–5273 (1979). 10.1073/pnas.76.10.5269

39. Danecek, P. et al. The variant call format and VCFtools. Bioinformatics 27, 2156–2158 (2011). 10.1093/bioinformatics/btr330

40. Peripolli, E. et al. Runs of homozygosity: current knowledge and applications in livestock. Animal Genetics 48, 255–271 (2017). 10.1111/age.12526

41. Purcell, S. et al. PLINK: a tool set for whole-genome association and population-based linkage analyses. Am. J. Hum. Genet. 81, 559–575 (2007). 10.1086/519795

42. Biscarini, F., Cozzi, P., Gaspa, G. & Marras, G. detectRUNS: an R package to detect runs of homozygosity and heterozygosity in diploid genomes. R package (2019). https://CRAN.R-project.org/package=detectRUNS

43. Mrode, R.A. Linear Models for the Prediction of Animal Breeding Values, 3rd edn. CABI, Wallingford, UK (2014). 10.1079/9781780643915.0000

44. Weir, B.S. & Cockerham, C.C. Estimating F-statistics for the analysis of population structure. Evolution 38, 1358–1370 (1984). 10.1111/j.1558-5646.1984.tb05657.x

