## Supplementary material for "Genomic signatures of long-term selection in the Langhill dairy herd: the power of a five-decade experiment for sustainable dairy cattle breeding": Suppl Notes

### Supplementary Note S1. Robustness of the XP-EHH and Rsb scans

#### Purpose

XP-EHH and Rsb depend on haplotype phase and on relatedness within lines, so we checked (i) that the published scores can be reproduced from the original inputs, (ii) how they depend on phasing and on the group of cows used, (iii) whether the signal is reproducible within the data, and (iv) whether the overlap between scans exceeds chance. Allele-frequency (ZFST) and diversity (π) scans do not use phase and are unaffected by (ii).

#### 1. Reproducing the published scans

The original inputs were the two separately phased files for the 1,073 concordant cows (496 Select, 577 Control). Re-running rehh with default settings on these files gave scores that correlated with the published values at r = 0.97 (XP-EHH) and r = 0.95 (Rsb) and placed the strongest peaks at the same positions (BTA10 22.3 Mb, 6.33 vs 6.11; BTA21 62.0 Mb, 5.80 vs 5.76; BTA12 71.6 Mb Rsb, 12.95 vs 9.90). The published scores were on a scale about 4% smaller than the rehh re-run, and fewer markers exceeded |4| (XP-EHH 70 vs 137; Rsb 177 vs 370), so the number of markers beyond a fixed threshold is sensitive to implementation details; positions and signs of the strongest signals were not. An independent implementation (scikit-allel XP-EHH) on the same files gave r = 0.91 with the published scores and the same two strongest peaks.

#### 2. Dependence on phasing and on the group of cows

| Phasing / cows (scikit-allel XP-EHH) | Markers \|z\|>4 | Max \|z\| | r with published | Top-0.1% regions also hit by ZFST/π |
| --- | --- | --- | --- | --- |
| 1 separate-phased, 1,073 (original groups) | 258 | 8.1 | 0.91 | 5/6 (chance 0.7, P=0.002) |
| 2 same, rehh-like (border markers dropped) | 258 | 8.09 | 0.91 | 5/6 (chance 0.7, P=0.002) |
| 3 jointly phased, 1,073 concordant | 2 | 4.04 | 0.35 | 0/10 (chance 1.3, P=1.000) |
| 4 jointly phased, all 1,126 recorded | 12 | 4.27 | 0.36 | 0/9 (chance 1.2, P=1.000) |
| 5 jointly phased, structure clusters 554/673 | 3 | 4.08 | 0.35 | 0/9 (chance 0.9, P=1.000) |
| N1 NULL: Control split in halves | 215 | 16.93 | 0.03 | 4/28 (chance 2.6, P=0.253) |
| N2 NULL: Select split in halves | 405 | 14.46 | 0.04 | 2/3 (chance 0.4, P=0.056) |

Only the separately phased groups reproduced the published scan. Re-phasing the same genotypes jointly across both lines (1,073 concordant cows, all 1,126 recorded cows, or the 554/673 structure clusters) gave 2–12 markers beyond |4| and no coincidence with ZFST/π regions, so the haplotype scans depend on within-line phasing. Random halves of a single line (rows N1, N2) produced 215 and 405 markers beyond |4| with heavy tails, so counts beyond a fixed threshold are not a test of selection.

#### 3. Reproducibility within the data

Random subsamples of 288 cows from the real Select-versus-Control contrast gave z-scores that correlated at r = 0.94–0.96 with the full run, and 57–75% of extreme markers recurred with the same sign (rehh re-run: recurrence 0.61, correlation 0.96). In random splits within one line the correlation between splits was close to zero (−0.13 to 0.03 in scikit-allel; 0.02 and −0.03 in the rehh re-run) and 7–9% of extremes recurred. A replicated core was defined as |z| > 4 in the full run, the same sign and |z| > 3 in every real subsample, and never |z| > 4 in any null split: 74 markers in 19 regions (scikit-allel) and 89 markers (rehh re-run). Of the 137 extreme XP-EHH markers in the rehh re-run, 100 were higher in the Select line and 37 in the Control line (scikit-allel: 237 of 254 higher in Select), consistent with the higher autozygosity and relatedness of the Select line.

| BTA | Start (bp) | End (bp) | Markers | Peak \|z\| |
| --- | --- | --- | --- | --- |
| 3 | 26,126,232 | 26,128,391 | 2 | 4.16 |
| 3 | 31,689,225 | 31,796,962 | 4 | 4.66 |
| 3 | 46,570,440 | 47,581,360 | 4 | 4.25 |
| 5 | 2,440,554 | 2,613,571 | 4 | 4.43 |
| 5 | 99,698,521 | 99,698,521 | 1 | 4.16 |
| 8 | 30,442,602 | 30,484,139 | 2 | 4.50 |
| 9 | 39,757,281 | 39,757,281 | 1 | 4.45 |
| 12 | 10,165,573 | 10,165,573 | 1 | 4.25 |
| 12 | 71,564,476 | 71,657,701 | 7 | 5.76 |
| 15 | 24,623,682 | 25,337,766 | 20 | 6.39 |
| 15 | 28,821,314 | 28,945,184 | 6 | 4.47 |
| 15 | 45,127,857 | 45,130,062 | 2 | 4.35 |
| 15 | 52,256,813 | 52,287,304 | 2 | 4.28 |
| 15 | 55,875,613 | 56,042,631 | 4 | 4.41 |
| 16 | 48,651,177 | 48,651,177 | 1 | 4.34 |
| 16 | 78,819,477 | 78,825,060 | 2 | 4.38 |
| 18 | 28,396,317 | 28,491,643 | 5 | 4.45 |
| 18 | 30,296,600 | 30,296,600 | 1 | 4.40 |
| 21 | 61,961,430 | 62,035,242 | 5 | 5.75 |

Table S1.1. Replicated core regions of the XP-EHH scan (scikit-allel run on the separately phased 1,073 concordant cows).

The BTA10 22.3 Mb peak, the highest published XP-EHH value, exceeded |z| = 5 in one of eight within-line null splits and is therefore treated as provisional.

#### 4. Convergence between scans compared with chance

Candidate regions from each scan were shifted at random along their chromosomes (1,000 replicates, lengths preserved). Regions supported by at least two scans numbered 45 (null mean 15, 95% interval 9–21), by at least three 17 (null 1, interval 0–3), and the single four-method region on BTA21 had P = 0.047. Shifting the two haplotype scans (XP-EHH and Rsb) together relative to the two frequency/diversity scans (ZFST and π), 41 of 84 haplotype regions coincided with a frequency/diversity region against 7.4 expected (P = 0.001). Because XP-EHH and Rsb are computed from the same phased haplotypes (15 of 34 top-0.1% locations shared within a run), they were not treated as independent.

Of the 17 regions supported by three or more scans, 14 include XP-EHH and 9 of these contain a replicated core region, including BTA21 59.2–62.2 Mb, BTA3 31.4–32.2 Mb, BTA8 29.9–30.5 Mb, BTA12 71.3–73.1 Mb, BTA15 24.8–25.4 Mb and BTA5 2.3–3.0 Mb. BTA21 and BTA3 are also supported by ZFST and π, which do not depend on phase, in every analysis variant.

#### 5. Summary

The published haplotype scans are reproducible from their original inputs; their signal is real in the sense that it replicates across random subsamples but absent from within-line splits; it exists only with within-line phasing; and its overlap with the allele-frequency and diversity scans exceeds chance. We therefore present XP-EHH and Rsb as complementary, phase-dependent evidence, and base the principal candidate regions (BTA21, BTA3) on convergence that includes phase-independent statistics.
